# Readout adaptation in simulated arm reaching: learned compensation and performance after updates stop

**DOI:** 10.64898/2026.06.11.731593

**Authors:** Jun Kobayashi

**Affiliations:** Department of Intelligent and Control Systems, Graduate School of Computer Science and Systems Engineering, Kyushu Institute of Technology, Iizuka, Fukuoka 820-8502, Japan

## Abstract

When an arm learns to compensate for a force that deflects its movement, removing the force can produce an opposite deviation, or after-effect. We evaluate what the controller retains by measuring the after-effect, the learned torque correction, and tracking performance after weight updates stop. We use a simulated torque-driven planar two-link arm reaching in eight directions under velocity-dependent sideways forces. Only an external linear torque readout adapts online; the parameters generating its features stay fixed. Features come from three configurations of closed-form continuous-time (CfC) networks, an echo-state network, or a radial-basis-function layer. All three CfC configurations, including neural circuit policy wiring, are trained for 400 epochs. These five feature-based variants use least-mean-squares (LMS) learning; a sixth uses recursive least squares (RLS) on the dense-backbone CfC readout. Learning rates are selected on three model initializations and evaluated on ten separate initializations under three force fields. All six variants improve tracking relative to their fixed-readout baselines and pass our after-effect criterion in every evaluation run. Diagnostics on the same ten evaluation initializations in the main field (60 conditions) quantify differences behind this shared decision: the median normalized squared residual between the learned output change and ideal compensation ranges from 0.0097 to 0.383 across implementations on common prescribed inputs. With updates stopped and feedback control active, all final readouts retain useful compensation, reducing tracking error by 40–97% relative to their pre-adaptation versions. Compared with the final adaptive cycle, the five LMS variants have median fixed-readout error ratios of 2.28–5.87, while RLS remains nearly unchanged. These measurements distinguish the performance achieved during adaptation from that retained in the final weights. They provide a concrete evaluation procedure for readout adaptation: measure the learned output correction and test the final controller with updates off, alongside behavioral after-effects. The comparison characterizes complete implementations in this task; differences in preparation and hyperparameters limit attribution to feature representations or update rules alone.

## 1 Introduction

In force-field reaching, an external force deflects the arm during a movement. Learning can reduce the trajectory error, and removing the force can then produce a deviation to the opposite side, an after-effect (Shadmehr & Mussa-Ivaldi, 1994). For an adaptive neural controller, an additional question is what the learned weights retain: the accuracy of the compensating output and the performance of the controller once further updates stop. We study these outcomes together, connecting a behavioral signature of adaptation to measurements of the learned output and the fixed final controller.

The setting is online adaptation of a linear torque readout on frozen features. This architecture connects cerebellar adaptive filters, reservoir readouts, and adaptive control with learned representations (Fujita, 1982; Dean et al., 2010; Sussillo & Abbott, 2009; O’Connell et al., 2022). We examine its use with offline-trained closed-form continuous-time (CfC) controllers (Hasani et al., 2022), including neural circuit policy (NCP) wiring (Lechner et al., 2020), and compare it with echo-state network (ESN) and radial-basis-function (RBF) features. CfC specifies the recurrent state update, while NCP specifies structured wiring. These implementations provide different fixed features to the same external readout. This setting is relevant both to researchers interpreting motor-adaptation models and to those assessing an online-adapted output layer for subsequent use with fixed weights.

Prior cerebellar controllers demonstrate force-field adaptation in simulation (DeWolf et al., 2016) and aftereffects on a physical robot (Casellato et al., 2014). Human experiments also show that after-effect magnitude depends on limb impedance as well as learned compensation (Osu et al., 2003). Our contribution is a quantitative comparison of three outcomes on the same model instances: catch-trial reversal, learned torque correction, and performance with the final readout fixed. A torque-driven planar two-link arm and a known curl force field allow us to compare the learned output change with the ideal field-compensating torque on common prescribed inputs. Fixed-readout tests then measure retained tracking performance in the closed loop, with PD feedback active in both the pre- and post-adaptation tests.

We compare three CfC configurations, ESN, and RBF using feedback-error learning with least-mean-squares (LMS) readout updates, plus recursive least squares (RLS) on the same dense-CfC features. All CfC configurations are trained for 400 epochs. Learning rates are selected on three model instances and evaluated on ten separate instances under three fields. The diagnostics cover the same 60 main-field conditions (six variants and ten instances); their questions were motivated by earlier exploratory results and their procedures fixed before this rerun. All six variants pass our sign-and-ratio after-effect criterion, while their learned torque corrections and fixed-final tracking errors differ. All final readouts retain useful compensation relative to their pre-adaptation weights; the five LMS variants have higher errors when fixed than during the last adaptive cycle, whereas RLS performs nearly the same. The resulting lesson is operational: assess what a readout has learned by measuring its output correction and testing its final weights with updates off, in addition to recording adaptation and after-effects.

## 2 Related work

### Linear adaptation on fixed features

Here, features are the numerical inputs to the linear output layer (readout), which forms a weighted sum of them to produce joint torques. In our controllers, these features are recurrent network states or RBF activations computed from the desired movement and current arm state. Fixed features means that the parameters generating them remain unchanged during online adaptation; the feature values themselves can vary over time. The adaptive-filter model of the cerebellum updates output weights on a fixed expansion with an error signal (Fujita, 1982; Dean et al., 2010), following the Marr–Albus architecture (Marr, 1969; Albus, 1971) and the LMS rule (Widrow & Lehr, 1990). In adaptive control, Sanner & Slotine (1992) adapt the output weights of a Gaussian RBF network with a stability-guaranteed law, and Neural-Fly (O’Connell et al., 2022) adapts linear coefficients on a learned, frozen representation online. Our RBF-LMS variant is structurally close to a fixed Gaussian-basis adaptive controller but implements plain LMS rather than a Lyapunov-based law, and our CfC readout adaptation in each configuration is the same idea applied to a frozen recurrent core.

### Continuous-time recurrent controllers

Liquid time-constant (LTC) and CfC models specify neural dynamics, whereas NCP specifies a structured sparse policy architecture (Hasani et al., 2021; 2022; Lechner et al., 2020). The Python library `ncps` supports NCP wiring with either LTC or CfC cells. All three CfC controller implementations in this study use CfC cells. These works train the full network offline; they do not study post-deployment readout adaptation, which is the setting here. We use their published model classes without modification.

### Reservoirs, readouts, and online recurrent learning

Reservoir computing separates fixed recurrent dynamics from a trained readout (Maass et al., 2002; Lukoševičius & Jaeger, 2009), and FORCE learning trains the readout online with RLS (Sussillo & Abbott, 2009). Cerebellar reservoir models make the same division (Yamazaki & Tanaka, 2007; Rössert et al., 2015). Online learning of recurrent weights themselves is possible, for example with random-feedback local online learning (Murray, 2019); we do not compare against such updates, and the frozen-core restriction is a design choice whose cost we do not measure.

### Force-field adaptation and after-effects

Human reaching adapts to velocity-dependent curl fields and shows after-effects on catch trials (Shadmehr & Mussa-Ivaldi, 1994); basis-function and multi-timescale models describe its generalization and dynamics (Thoroughman & Shadmehr, 2000; Donchin et al., 2003; Smith et al., 2006), and internal-model and FEL accounts link it to cerebellar learning (Wolpert et al., 1995; Kawato, 1999; Gomi & Kawato, 1993; Tseng et al., 2007). Human force-field studies have already shown that after-effect magnitude reflects both predictive compensation and limb impedance (Osu et al., 2003); error-clamp experiments provide additional measurements of retained adaptation (Smith et al., 2006). Spiking neural controllers with cerebellar components demonstrate force-field adaptation in simulation (DeWolf et al., 2016). Casellato et al. (2014) use after-effects during field removal as evidence of predictive compensation in robotic reaching. We complement these behavioral observations by measuring the learned torque correction and testing the final readout with updates stopped. Our sign-and-ratio criterion is one specified summary of catch-trial behavior, evaluated alongside these other measurements.

## 3 Methods

### 3.1 Arm, trajectories, and protocol

Figure 1 summarizes the control and learning paths. At each time step, the feature generator receives the desired movement and measured arm state. Its external readout supplies one torque command, and the PD controller supplies a correction based on position and velocity errors. Let *z*_*t*_ denote the feature vector and *J*_*t*_ the readout weight matrix. The readout and PD torques sum to the commanded joint torque, 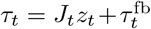. The external force acts separately on the arm. During adaptation, the PD torque also supplies the readout learning signal. Fixed-readout tests retain both the readout output and PD feedback while disabling learning.

**Figure 1:**
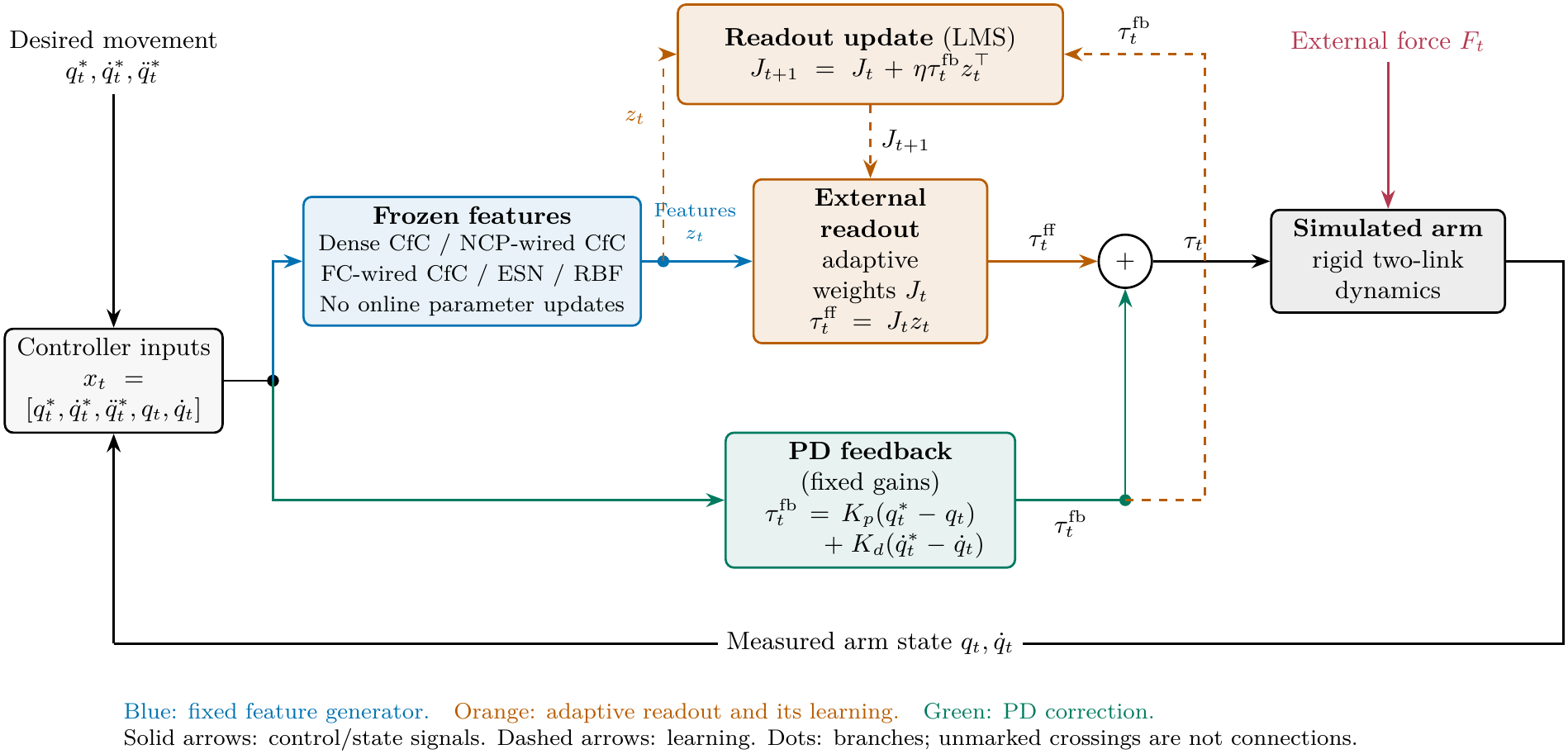
Control architecture and online readout learning. A fixed feature generator supplies features to an external linear readout. The readout torque and PD feedback torque are summed to drive the simulated arm, which is also subject to an external force. During adaptation, the features and PD torque are used to update the readout weights; the feature-generator parameters remain fixed. Solid arrows show control and state signals, and dashed arrows show learning.

The diagram depicts the five feature-based variants with LMS readout adaptation: dense CfC, NCP-wired CfC, fully connected wired CfC, ESN, and RBF. Dense-CfC RLS uses the same control paths with a different readout update rule.

The plant is a horizontal two-link arm with no gravity, contact, or actuator limits. Its dynamics are

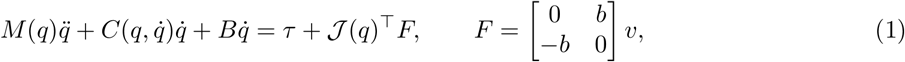

where *q* ∈ R^2^ is the joint configuration and *v* is end-effector velocity. The field strength *b* is measured in N s m^−1^. The arm is a rigid two-link linkage driven by joint torques, with link lengths 0.30 and 0.33 m, link masses 1.6 and 1.1 kg, center-of-mass offsets 0.11 and 0.16 m, inertias 0.025 and 0.045 kg m^2^ about the centers of mass, and joint viscous damping 0.05 N m s rad^−1^; it has no muscles, actuator dynamics, or sensor noise. A fourth-order Runge–Kutta solver uses a 5 ms step. Each reach starts at (0, 0.45 m) and ends at one of eight targets equally spaced on a circle of radius 0.10 m around that starting point. The targets are visited in a fixed order. Each reach lasts 0.5 s and follows a minimum-jerk trajectory. Feedback torque is

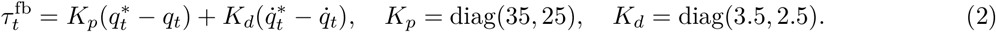

Each cycle consists of eight reaches, one to each target. Each run begins with one baseline cycle without an external force or readout updates (phase A). This is followed by 30 cycles with the force field present and readout adaptation enabled (phase B). The force field is then removed for one catch cycle, with readout updates disabled, to measure the after-effect. Finally, five washout cycles are performed without the force field and with readout adaptation enabled. The recurrent state resets at each reach boundary while the adaptive readout persists. During adaptation, we first compute the current torque command using the current readout weights. We then update those weights using the PD feedback torque as the learning signal, 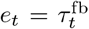, following feedback-error learning. This signal represents the correction supplied by the feedback controller, rather than a directly measured difference between the ideal torque and the readout output. The arm is advanced using the already computed torque command, so the updated weights affect outputs from the next time step onward. This update timing is an algorithmic choice, and the learning scheme is a design-level analogy to cerebellar error-driven learning, not a circuit-level model (Gomi & Kawato, 1993; Dean et al., 2010).

### 3.2 Controller preparation and online readout adaptation

The ten-dimensional controller input is 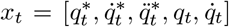. During offline training without an external force field, the measured-state entries *q*_*t*_ and 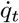 are replaced by the desired values 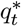 and 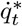 (teacher forcing). The target output is the arm’s analytic null-field inverse-dynamics torque. The dataset contains 24 sequences: eight directions at each of three radii, 0.08, 0.10, 0.12 m. These are split into 18 training and six validation sequences separately for each seed, with the same split used across the three CfC configurations within that seed.

Figure 2 compares the internal structures of the three CfC controllers and their common external torque readout. The controller in Figure 2(a) uses the standard CfC cell (Hasani et al., 2022) implemented in ncps, a Python library for continuous-time recurrent networks and structured neural wiring.^1^ Inside this cell, a fully connected layer with 128 units computes intermediate features for the recurrent state update. We refer to this configuration as dense-backbone CfC, following the term “backbone” used by Hasani et al. (2022) for the internal feature-processing network. Its external torque readout receives the 64 recurrent states, rather than the 128 backbone activations.

**Figure 2:**
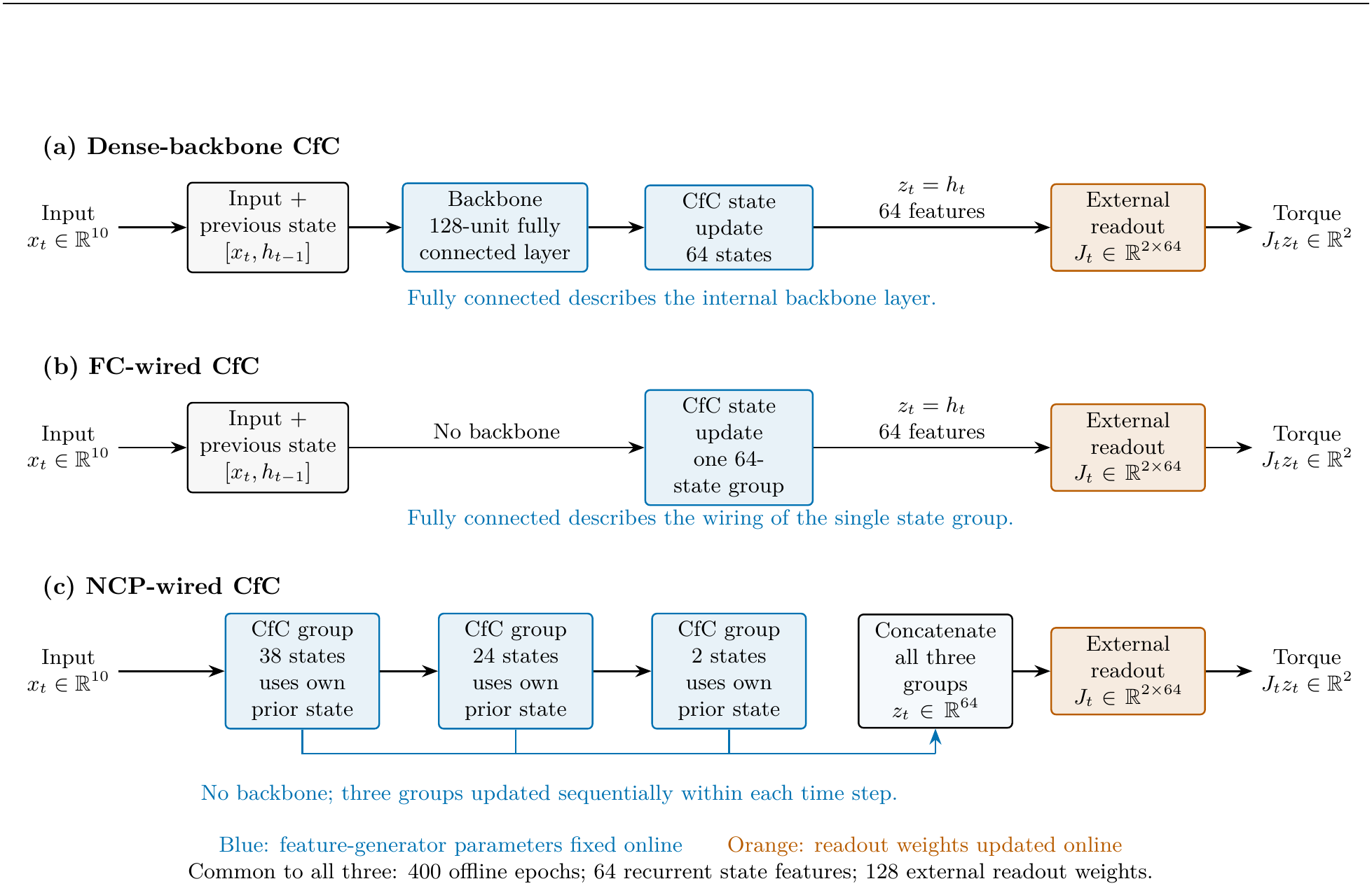
Three CfC controller configurations. The configurations differ in their internal backbone and recurrent-state grouping, but all supply 64 state features to an external linear torque readout. Blue blocks remain fixed during online adaptation; orange readouts adapt. Arrows show computation paths, not individual synaptic connections.

The controller in Figure 2(b) has a single fully connected group of 64 recurrent states and no backbone. Its wiring is specified using the `ncps class FullyConnected` (64, 2), where the arguments specify 64 state neurons and two designated output neurons. The external torque readout uses all 64 states. We call this controller FC-wired CfC. Here, fully connected refers to the wiring of the state group, rather than an additional internal layer. The state-update parameterization therefore differs from that of dense-backbone CfC.

The controller in Figure 2(c) has three sequentially updated groups of 38, 24, and 2 recurrent states and no backbone. Its wiring is specified using the `ncps` class `AutoNCP(64, 2)` with sparsity setting 0.5. As in the fully connected wiring, the arguments specify 64 state neurons and two designated output neurons, called motor neurons in NCP wiring. We call this controller NCP-wired CfC. The external readout receives the concatenated states of all three groups, rather than only the two motor neurons. Fixed connectivity masks generated by `AutoNCP` preserve selected connection restrictions during training. In the installed implementation, these restrictions apply to candidate-output weight matrices; within-group recurrent blocks and time-gating matrices remain dense. Appendix C gives the mask and implementation-audit details.

All three configurations supply 64 recurrent state features to the external readout. In compact labels, Dense, FC, and NCP denote dense-backbone, FC-wired, and NCP-wired CfC, respectively. The core trainable parameter counts are 42,624 for Dense, 19,200 for FC, and 13,712 for NCP; the external readout adds 128 parameters in each case. These configurations differ in parameterization and group structure as well as connectivity. FC-wired and NCP-wired CfC differ in both connectivity and state grouping, so their performance difference cannot be attributed to connectivity alone.

All three CfC configurations are trained for 400 epochs using full-batch Adam with learning rate 10^−3^ and the final-epoch checkpoint. All thirteen controller seeds (three selection seeds and ten evaluation seeds; Section 3.3) are prepared by the same configuration-specific procedure, with no model excluded based on fit. NCP wiring seeds are 22300–22302 for selection seeds 100–102 and 22310–22319 for evaluation seeds 110–119. Thus NCP variability includes wiring variability. The preliminary experiment used wiring seed 22222 throughout.

For comparison with CfC features, we use two alternative feature generators: an ESN, whose recurrent weights remain fixed, and an RBF layer, whose activations depend on the distance between the input and fixed centers. The ESN provides 64 recurrent features from a leaky tanh reservoir with standardized inputs.

Input weights are uniform on [−*s*_in_, *s*_in_], recurrent weights are Gaussian and scaled to spectral radius *ρ*, and biases are uniform on [−0.1, 0.1]. Its state starts at zero for each reach. The grids for spectral radius *ρ*, input scale *s*_in_, and leak *a* are {0.5, 0.9, 1.2} × {0.1, 0.3, 1} × {0.1, 0.3, 1}.

The RBF layer provides 64 features from Gaussian basis functions with k-means centers fit on standardized training inputs. Its Gaussian width is a multiplier times the median nearest-center distance, with candidate multipliers {0.5, 1, 2, 4, 8}. Each ESN/RBF readout is initialized by ridge regression with *λ* ∈ {10^−4^, 10^−3^, 10^−2^, 10^−1^, 1}. The ridge penalty is 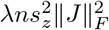, where *n* is the training sample count and 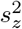 is the mean feature variance; this normalizes one aspect of feature scale across bases.

The ESN reservoir settings and RBF width multiplier are selected on seeds 100, 101, 102 by mean validation normalized mean-square error; the selected settings are then instantiated for all seeds. For each seed, its ridge penalty is chosen on that seed’s validation split, as specified before online evaluation. The chosen ESN setting is (*ρ, s*_in_, *a*) = (0.5, 0.3, 1); the selected *a* = 1 makes the leaky update a standard tanh recurrence. The chosen RBF width multiplier is 8, the upper edge of its predeclared grid.

During online adaptation, the dense-backbone, FC-wired, and NCP-wired CfC, ESN, and RBF readouts all use the same LMS form:

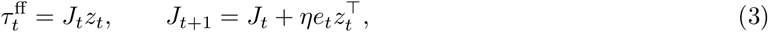

with *J ∈* R^2×64^ and without weight decay in the online readout update. To compare readout learning rules using the same features, we also update the dense-backbone CfC readout with regularized RLS, using a fixed forgetting factor *β* = 1, and *P*_0_ = 0.1*I* (Sussillo & Abbott, 2009); its parameters are not tuned in this comparison. The fixed-readout baselines keep each model’s initial readout *J*_0_ unchanged throughout the run. The PD-only baseline uses feedback torque alone, with no feedforward torque.

### 3.3 Predeclared selection, evaluation, and metrics

For each model family using LMS, we test nine candidate learning rates on three independently generated model instances, identified by random seeds *S* = {100, 101, 102}. We select the candidate with the lowest mean final exposure error among those satisfying the validity criteria given below on all three instances. The learning rate is therefore selected by measured performance, not drawn at random. We then hold this rate fixed and evaluate it on ten new model instances (*E* = {110, …, 119}), without using their outcomes to revise the selection. All model families use the same arm dynamics, direction order, force fields, and trial schedule; using the same seed number across families does not imply identical random draws.

We scale the learning-rate candidates by the average squared magnitude of each model’s feature vector. For model family *f*, let *g*_*f*_ be the mean squared Euclidean norm 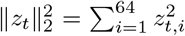, averaged over the null-field training samples within each selection seed and then over the three selection seeds. Let *g*_ref_ denote this value for dense-backbone CfC. We use nine base learning rates *α*: 0.00003, 0.0001, 0.0003, 0.001, 0.003, 0.01, 0.03, 0.1, and 0.3. Each base rate is scaled for model family *f* to give the candidate learning rate

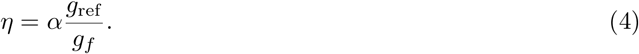

This scaling assigns smaller learning-rate candidates to models with larger average squared feature norms. It changes the candidate rates, not the LMS update itself, and does not match actual online update norms, feature covariance, or learning dynamics. Grid values were fixed before evaluation. The selected rates are dense CfC 0.0100, NCP-wired CfC 0.01625, FC-wired CfC 0.01102, ESN 0.04823, and RBF 0.001168. None lies at an online-grid edge. The wider grid was fixed for this rerun after inspecting a preliminary experiment; it was not expanded after observing these evaluation outcomes.

Learning rates are selected under a velocity-dependent curl field with strength *b* = 13 N s*/*m. A candidate is valid only if the initial null-field cycle and all 30 field-exposure cycles are complete, all reaches have finite numerical values, the final exposure-cycle error is lower than the first exposure-cycle error, and no exposure-cycle error exceeds twice the first exposure-cycle error. Among candidates valid for all three selection seeds, we choose the rate with the lowest mean final exposure-cycle error, with smaller *η* breaking ties. Catch and washout outcomes do not enter selection. The selected learning rates are then evaluated on ten independent model instances under three curl-field conditions: *b* = −13 N s*/*m, used for rate selection; *b* = 13 N s*/*m, with the force direction reversed; and *b* = 25 N s*/*m, with a stronger field. For each model family, seed, and force field, we also run a baseline that keeps the readout weights fixed at their initial values throughout the experiment (B0). Each adaptive model is compared with its corresponding baseline. The machine-readable results additionally report the dense-backbone CfC baseline alongside results from the other model families. No significance test of family superiority is applied to these ten descriptive evaluation seeds. For the primary dense/NCP comparison at *b* = 13, we report all seed-pair values and a 95% percentile bootstrap interval for the mean log error ratio, using 10,000 paired resamples and a fixed bootstrap seed. Pairs share the data split, not identical weights or wiring. The interval is descriptive and does not establish equivalence or a single-factor architecture effect. Non-finite or non-positive pairs would be listed separately; none occurs in this comparison. The machine-readable per-seed evaluation table records A null error, *B*_first_, *B*_last_, catch, washout, both B0 references, direction-level mirror counts, finite-run flags, and the selected learning rate for every method–seed–field combination (Appendix A).

The primary trajectory metric is the cycle mean of the maximum absolute perpendicular deviation, in millimeters. *B*_first_ is measured after the first exposure cycle, during which adaptation already occurs; *B*_last_ is the error in the final exposure cycle with adaptation. Improvement relative to the fixed-readout baseline is 1 − (*B*_last_*/B*_0,last_), where *B*_0,last_ is the final exposure-cycle error in the fixed-readout baseline (B0) for the same model family, seed, and force field. Figure 3 illustrates the signed deviations used to assess catch-trial reversal. Let *c*_*d*_ denote the catch-trial signed deviation and *r*_*d*_ the reference signed deviation from the final field-exposure cycle of the fixed-readout baseline (B0), for reach direction *d*. Their averages over the eight reach directions are 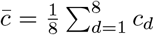 and *r*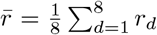. For 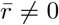, the after-effect ratio and mean-direction mirror flag are

**Figure 3:**
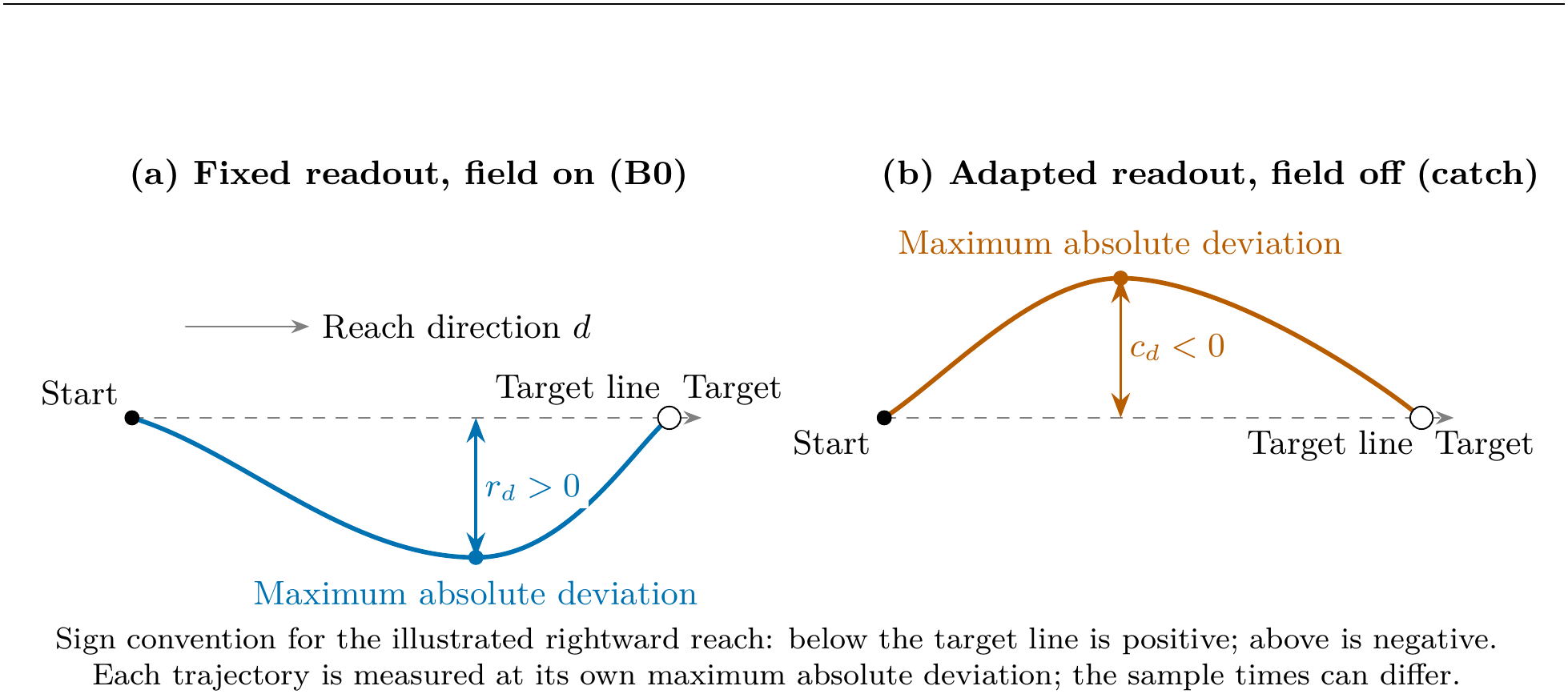
Measurement of *r*_*d*_ and *c*_*d*_ for one reach direction (schematic, not measured trajectories). Each is the signed perpendicular distance from the target line at the trajectory’s maximum absolute deviation: (a) the final field-exposure cycle of the fixed-readout baseline and (b) the field-free catch cycle after adaptation. The illustrated deviations have opposite signs.

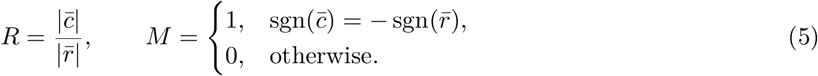

Each signed direction value is the signed deviation at the time of maximum absolute perpendicular deviation, which can occur at different times across directions. If *r*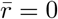, both the ratio and the mirror flag are treated as missing. The prespecified adjudication requires *M* = 1 and *R*≥ 0.25; we also report the number of individual directions with opposite signs and thresholds 0.10, 0.15, 0.20, 0.25, 0.30, 0.40, 0.50 . The adaptation-speed indicator is the first B cycle attaining half the reduction from *B*_first_ to *B*_last_, counted from one. Washout decay is the magnitude of the fifth washout-cycle signed deviation divided by catch magnitude; it is missing if catch magnitude is below 1 mm. Non-finite cycles contribute missing values, not zero. Recorded readout norms are cycle-end values from the last reach of each cycle, not maxima over all time steps.

In a sensitivity analysis, we multiply both PD gain matrices, *K*_*p*_ and *K*_*d*_, by the same factor (0.5, 1, or 2) and test all nine candidate learning rates for each of the three CfC configurations, ESN, and RBF, using only the three model instances used for learning-rate selection. Family-specific B0 runs at each gain supply the after-effect reference. The × 1 adaptive runs are identical to and reused from selection; these outcomes are descriptive and do not revise the selected learning rates.

### 3.4 Exploratory compensation and fixed-readout diagnostics

Before the 400-epoch rerun, exploratory simulations on earlier model instances (seeds 0–2) compared learned torque corrections with ideal field compensation and tested tracking with the final readout weights held fixed. Those results motivated the diagnostic questions examined here. The diagnostic procedures were specified before the 400-epoch online rerun, and were then applied at *b* = 13 to six methods on each of the ten new evaluation model instances (seeds 110–119; 60 conditions). Neither torque-probe nor fixed-readout outcomes enter learning-rate selection. Although the evaluation instances are new, these diagnostics follow up prior observations rather than constitute a preregistered confirmatory study.

Let *J*_pre_ be the offline readout and *J*_post_ the readout after 30 exposure cycles, before catch. We regenerate the same adaptive protocols without washout to save these states; catch itself has adaptation off. For a fixed nominal reach, the controller receives 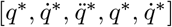, with hidden state reset per reach. The same features drive both frozen readouts. Their output difference *δτ* is compared with

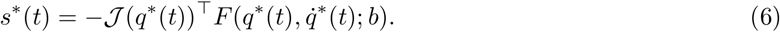

For two torque signals *u* and *v*, we define the inner product as

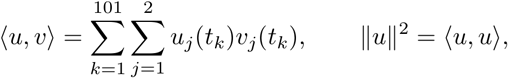

where *t*_*k*_ indexes the 101 controller evaluation times and *j* indexes the two joints. All times and joints have equal weight. Thus *D* = ⟨*s*^*\**^, *s*^*\**^⟩ is the sum of squared ideal compensation torques over these times and joints. Using *D* for normalization, we report

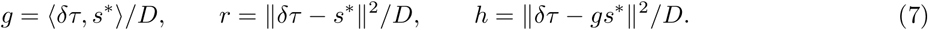

These are a projection coefficient and normalized squared residuals, not percentage torque errors. We verify *r* = (*g* − 1)^2^ + *h*. The best time-constant torque is 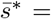 mean_*t*_ *s*^*\**^, giving residual 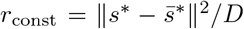. Only *r < r*_const_ implies a closer match than the best constant on that input; *h* is unsuitable because it is zero even for zero output change. We also record the pre-readout null-dynamics error, total post-readout target error, and their cross term. A small baseline error does not causally separate baseline correction from field learning.

For the closed-loop test, each of the 60 pre/post readout pairs is run with updates off at *b* = 13, using the same eight trained reaches, initial states, and PD gains: 960 reaches, with no further learning. We also archive common-input readout comparisons on the realized fixed-readout trajectories. These supplementary probes are defined in Appendix C, with values provided in the accompanying data.

We compare fixed-readout tracking error with the corresponding B0 and adaptive first/final cycles. The adaptive final cycle has changing weights across time and directions; the fixed test applies its single final readout to every direction. Their difference does not isolate dependence on within-reach learning. Feedback remains active in every test. The fixed-post reduction is 1 − (*E*_post_*/E*_pre_), not the fraction of adaptive error reduction retained. The tests assess the same trained movements, not generalization to new movements.

## 4 Results

### 4.1 Online adaptation on separate evaluation seeds

Table 1 summarizes final tracking performance and catch-trial deviations in the main field (*b* = 13 N s*/*m). Figure 4 shows the exposure-error trajectories and individual model instances’ signed catch deviations under the same conditions.

**Table 1:** Online adaptation at *b* = 13 on evaluation seeds 110–119. First B, Final B, Reduction, and Catch are medians across ten seeds. AE counts passing seeds; Reversed /8 gives the minimum–maximum direction count across seeds. Reduction uses each model’s own frozen-readout reference. The after-effect decision uses the sign and direction-mean ratio threshold 0.25. Direction counts describe target directions within a run, not independent samples.

| Method | First B (mm) | Final B (mm) | Reduction (%) | Catch (mm) | AE | Reversed /8 |
|---|---|---|---|---|---|---|
| Dense CfC LMS | 5.60 | 0.856 | 96.4 | -22.85 | 10/10 | 8 |
| NCP-wired CfC LMS | 5.57 | 0.906 | 96.2 | -21.85 | 10/10 | 8 |
| FC-wired CfC LMS | 6.25 | 0.773 | 96.8 | -22.68 | 10/10 | 8 |
| ESN LMS | 2.19 | 0.188 | 99.2 | -18.89 | 10/10 | 8 |
| RBF LMS | 9.92 | 5.161 | 78.4 | -11.51 | 10/10 | 7 |
| Dense CfC RLS | 5.13 | 0.793 | 96.7 | -23.82 | 10/10 | 8 |

**Figure 4:**
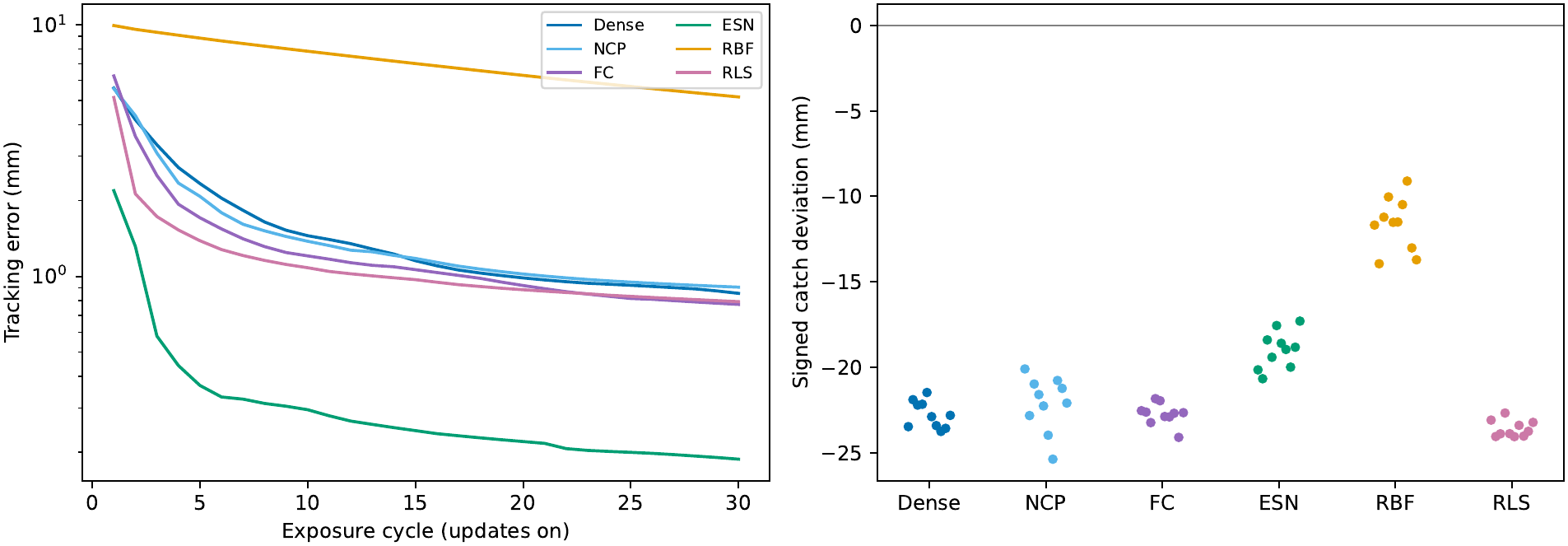
Online adaptation on evaluation seeds 110–119, *b* = 13. Left: median exposure error with updates on. Right: individual signed catch deviations. Dense, NCP, and FC identify the three CfC configurations. Every adaptive run passes the direction-mean catch criterion; RBF need not reverse every individual direction.

At *b* = −13, dense and NCP-wired CfC have median final adaptive errors of 0.856 and 0.906 mm, versus approximately 24 mm for their frozen-readout references. The median family-referenced reductions are 96.4% and 96.2% (Table 1). The geometric mean NCP/dense error ratio is 1.131 across ten paired seeds; the mean log ratio is 0.123 with descriptive bootstrap interval [0.015, 0.258] (ratio scale approximately [0.985, 1.294]). Dense has smaller error in eight pairs and NCP in two. This comparison neither establishes equivalence nor identifies a generally better configuration.

FC-wired CfC, ESN, RBF, and RLS also reduce tracking error relative to their own references. Selected rates are reused at *b* = −13 and *b* = 25 without retuning. All 180 adaptive evaluation runs are finite and pass the sign-and-ratio criterion at threshold 0.25. The five non-RBF methods reverse all eight directions. RBF reverses seven directions in all ten main-field runs, and in 29 of its 30 runs over all fields. In all ten main-field runs, its non-reversed direction is the fifth reach direction (index 4), which also has the smallest median absolute deviation in the corresponding fixed-readout baseline (5.93 mm). These outcomes establish bounded readout recalibration but not specificity of the coarse catch decision.

### 4.2 Learned compensation on common inputs

Although all 60 main-field conditions pass the same catch-trial criterion, their nominal compensation residuals differ across implementations and model instances (Table 2; Figure 5). These are normalized squared residuals on common prescribed inputs, not percentage torque errors or an isolated ranking of feature bases. Among the five variants that reverse all eight directions, ESN has both the smallest median nominal compensation residual and the smallest median after-effect ratio (0.80; median signed catch deviation −18.9 mm).

**Table 2:**
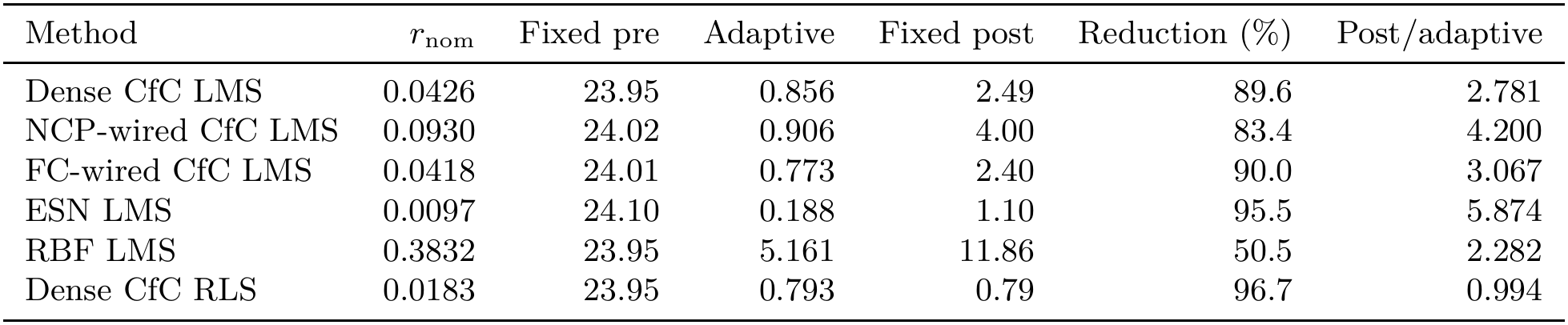
Compensation and fixed-readout diagnostics on the same ten evaluation seeds, *b* = 13. Each entry is a median over ten runs; *r*_nom_ first takes a median over the eight directions within each run. Tracking errors are in millimeters. Pre/post refer to readouts before/after exposure, held fixed with the same active PD feedback. Reduction is 100(1 −*E*_post_*/E*_pre_); post/adaptive is computed within each seed before taking its median. All conditions pass the coarse catch decision, but RBF reverses seven of eight directions.

**Figure 5:**
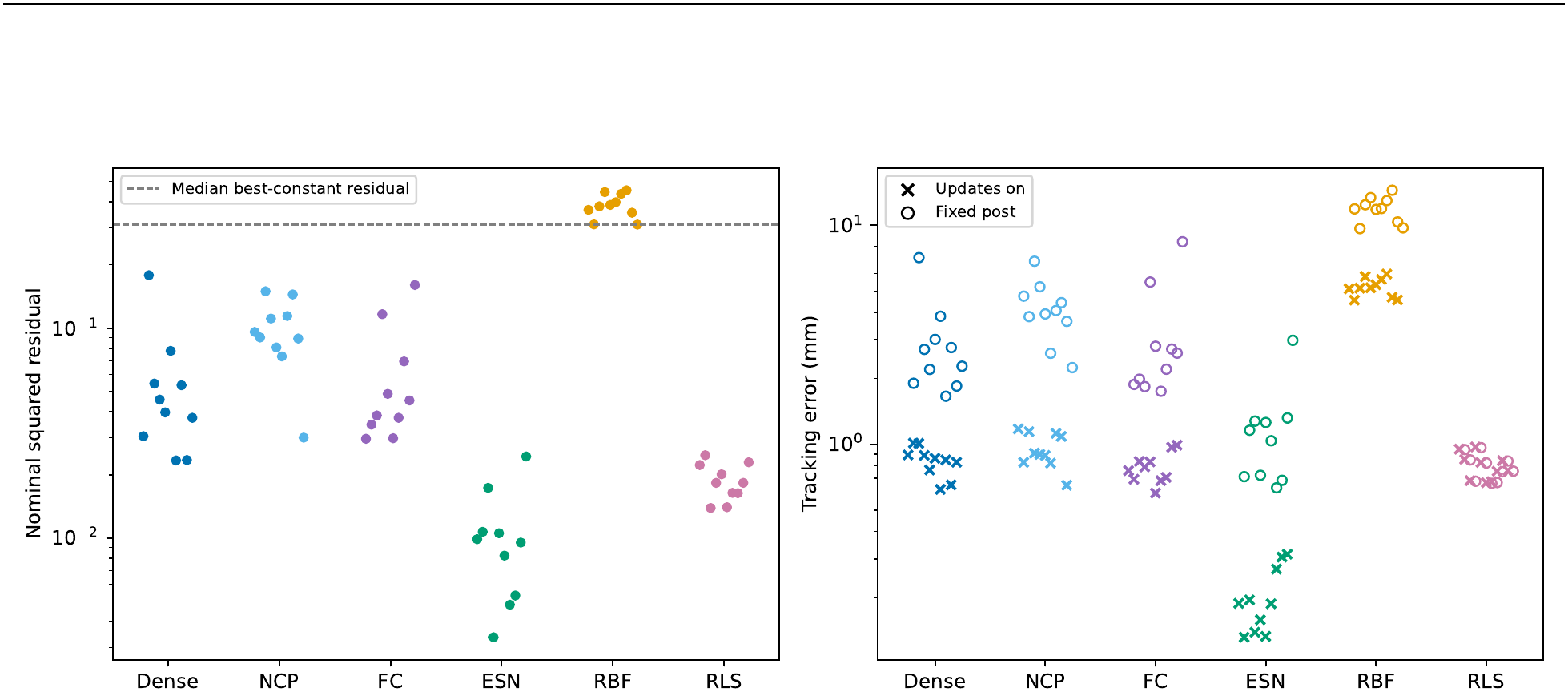
The same 60 evaluation-seed conditions, *b* = 13. Left: each run’s direction-median nominal compensation residual; the dashed line is the median best-constant residual. Right: adaptive final-cycle errors (crosses) and fixed-post errors (open circles), with identical PD gains. Each point is a model individual; its position within a group is only for separation. Absolute errors accompany ratios in Table 2.

Applying the same direction-then-seed median aggregation on inputs from the realized fixed-post trajectories gives residuals of 0.0415 (dense CfC LMS), 0.0056 (ESN), 0.385 (RBF), and 0.0172 (dense-CfC RLS), preserving the broad pattern of the nominal-input results (Appendix C; accompanying data).

The median best-constant residual across directions is 0.311. Dense and FC-wired CfC achieve smaller residuals than the best per-direction constant correction in 78 of 80 seed–direction observations, NCP in 73, ESN and RLS in all 80, and RBF in 27. Directions are nested within ten model individuals and are not independent replications. Learned output waveforms for a fixed illustrative seed, direction, and joint are shown in Figure 6. The full arrays include every direction and both joints.

**Figure 6:**
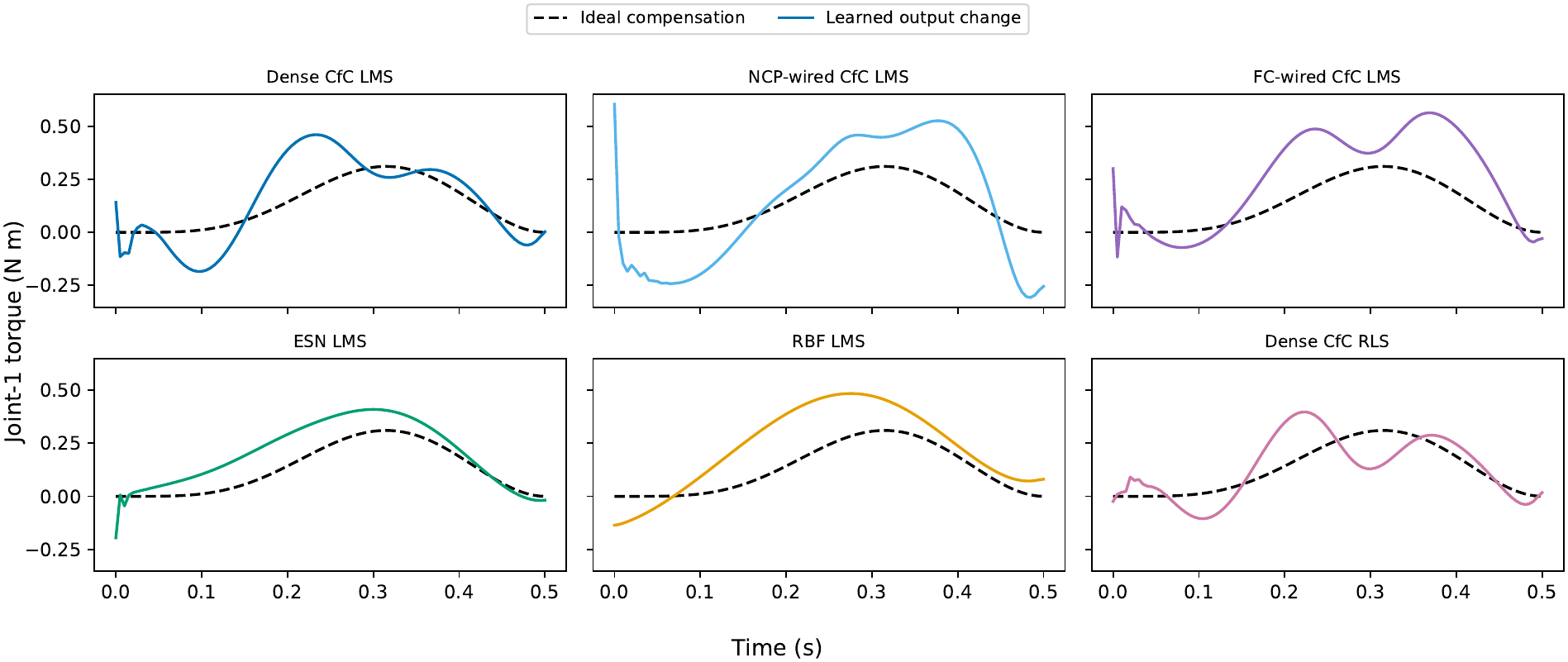
Nominal learned output change and ideal field compensation, evaluation seed 110, direction index 0 (the first target), and the first joint (shoulder). The same illustrative index is used for every method. All controller sample times, including the initial transient, are retained.

LMS and RLS use identical dense-CfC features within a seed. Their different attained residuals show that an LMS outcome cannot be treated as the best representation achievable by that feature basis. This comparison does not isolate the update rule from its hyperparameters or solve for an optimal readout. The 50 non-RBF conditions all reverse every direction and still exhibit different residuals; this restricted observation is descriptive.

### 4.3 Performance with the final readout fixed

All 60 final readouts reduce field-on tracking error relative to their pre-exposure readouts when updates stop and PD feedback remains active. Reductions range from 39.9% to 97.3%. The ten-seed median reductions are 89.6% (dense), 83.4% (NCP), 90.0% (FC), 95.5% (ESN), 50.5% (RBF), and 96.7% (RLS). These compare fixed post with fixed pre; they are not the fraction of adaptive improvement retained.

Compared with the final cycle with updates on, fixed-post errors are larger for all five feature-based LMS variants in every evaluation seed. Their median post/adaptive ratios are 2.78, 4.20, 3.07, 5.87, and 2.28 for dense, NCP, FC, ESN, and RBF, respectively. RLS has ratios of 0.991–0.995, so its fixed-final performance is nearly unchanged under these settings. Median absolute fixed-post errors are 2.49, 4.00, 2.40, 1.10, 11.9, and 0.79 mm in table order. A large post/adaptive ratio can accompany a small absolute error, and the ratio is not evidence of a particular mechanism.

Direction-level reanalysis shows that the seed-median fixed-post/adaptive error ratio exceeds one in all eight directions for every LMS variant, including the last direction in the training order (medians 2.15–2.80 across the five LMS variants). Some individual seed–direction pairs improve when fixed. Thus the median increase extends across the trained directions; it is not confined to directions preceding the last one. The accompanying data provide the paired direction-level errors and ratios.

### 4.4 Sensitivity of the catch criterion

The selection-seed rate/gain grid contains 405 adaptive outcomes, of which 349 are finite. Six finite runs pass the catch criterion while final error exceeds their own gain-matched frozen reference: four large-rate RBF conditions and two low-rate dense/FC conditions. All six fail the learning-rate selection validity rules and therefore are not eligible for the selected evaluation comparison. They illustrate the behavior of the catch criterion when used alone, rather than the combined tracking and validity checks used to select learning rates. For the four RBF cases, final errors are approximately 307–585 mm. Exceeding the fixed-readout reference does not imply that error never decreased after the first adaptive cycle.

Threshold sensitivity is also not invariant: all 180 evaluation runs pass thresholds 0.10 through 0.30, 179 pass at 0.40, and 163 pass at 0.50 (the exceptions are RBF). Appendix B reports the gain/rate panel, including invalid points.

## 5 Discussion

The combined measurements quantify what is retained by an online-adapted readout. On the same model instances, a common catch-trial decision accompanies different torque corrections, and useful fixed-final performance coexists with a gap relative to ongoing adaptation. For researchers evaluating motor-adaptation models, this supplies an output-level check alongside the behavioral signature. For those using adapted readouts with learning disabled, it supplies a direct test of the controller that will actually be used. The final-weight test adds information even when the adaptive learning curve has reached a low error.

Adaptive-control theory distinguishes tracking convergence from parameter convergence, with parameter identification depending on sufficient excitation of the regressor (Boyd & Sastry, 1986). A repeated reaching schedule alone does not establish that condition for the 64-dimensional feature vector. Adaptive-filter theory also distinguishes constant-step LMS, whose weights can continue fluctuating under stationary stochastic inputs (Widrow et al., 1976), from the covariance-dependent gains of no-forgetting RLS, which can decrease as observations accumulate. In FORCE learning, RLS can establish static weights that sustain the target after learning stops (Sussillo & Abbott, 2009). These results motivate testing retained performance rather than inferring it from adaptive tracking. Our deterministic closed-loop feedback-error learning experiment is not the stochastic filtering or model-reference setting of those analyses. The direction-level results constrain the observed gap: its median extends to all trained directions, including the last. They do not distinguish withinreach updates, parameter variation across directions, or incomplete convergence as causes, nor establish a batch-optimal RLS solution.

The three CfC implementations demonstrate readout recalibration after equal numbers of offline training epochs, across ten separate evaluation initializations. The comparison with ESN and RBF broadens the set of fixed features exhibiting retained compensation; the dense-CfC LMS/RLS comparison shows how the attained correction and fixed-final performance can differ even on the same features. These are comparisons of complete implementations: backbone structure, recurrent grouping, effective connectivity, parameter count, offline fit, initialization, wiring, and hyperparameter settings all contribute. FC-wired CfC provides an additional implementation comparison rather than a one-factor wiring ablation. The dense/NCP paired result remains uncertain.

The learning-rate search selected interior grid values for all five tuned variants, while RBF width remained at its offline grid edge. Feature-norm scaling adjusts the rate candidates using one offline scalar; further matching of update magnitudes or feature covariance would be needed to attribute the performance differences to representation alone. The cerebellar interpretation likewise concerns the computational design of a fixed feature map, plastic output weights, and a feedback-based learning signal (Fujita, 1982; Dean et al., 2010). The present measurements extend the evaluation of such controllers at the torque and fixed-controller levels; comparison with human data would be a separate test of biological adequacy.

## 6 Limitations

All results concern a torque-controlled horizontal two-link arm without gravity, contact, actuator limits, three-dimensional motion, or physical validation. Evaluation and diagnostics use ten model initializations, not ten human subjects. Diagnostic questions and the wider grid were informed by earlier results, then fixed before this rerun; the study is not a preregistered confirmatory test. Fixed tests reuse trained directions, radius, speed, plant, and PD gains and do not establish movement generalization.

Matching the CfC epoch count does not match compute, capacity, or offline fit, nor does it make the other feature generators equally trained. NCP wiring changes across individuals. No recurrent-core plasticity, RFLO, or matched adaptive-control law is tested. Descriptive intervals over ten shared-split pairs do not establish equivalence or population-wide superiority.

Nominal probes evaluate functions on chosen inputs. Realized-state probes are at controller sample times, not an energy balance over RK4 substeps or a decomposition of feedforward and feedback effects. No new cross-controller A/catch input-replay study is included in this rerun. The direction-level data constrain the distribution of the adaptive/fixed gap but do not decompose its causes or measure convergence of the readout weights.

## 7 Conclusion

In this simulated arm-reaching task, readout adaptation leaves useful compensation in the final weights of all six tested variants. The learned correction and fixed-final performance distinguish these implementations even though every adaptive evaluation run passes the same after-effect criterion. With feedback active, all 60 final readouts improve tracking relative to their pre-adaptation versions. The five LMS variants show higher errors with the final weights fixed than during the last adaptive cycle, while RLS remains nearly unchanged.

The practical contribution is a joint evaluation of behavior, learned output, and retained controller performance on the same model instances. For readout-adaptation models, this links a behavioral signature to the content of the learned output; for controllers used after adaptation, it measures the performance available from the retained weights.

## A Per-seed main-field outcomes

Table 3 reports all 60 adaptive main-field evaluation rows. The machine-readable dataset also includes the family-specific fixed-readout baselines, PD-only, offline fit, both after-effect references, finite flags, and the other fields. PD-only has no learned model or offline validation error; any computational reference checkpoint is only an implementation convenience.

**Table 3:** Main-field individual adaptive outcomes, evaluation seeds 110–119. Tracking errors and catch deviations are in millimeters. Offline fit and both reference baselines are included in the machine-readable supplement.

| Method | Seed | First B | Final B | Catch | AE ratio | Reversed /8 |
|---|---|---|---|---|---|---|
| Dense | 110 | 5.78 | 0.894 | -23.47 | 0.984 | 8 |
| Dense | 111 | 5.55 | 1.012 | -21.90 | 0.913 | 8 |
| Dense | 112 | 6.50 | 1.010 | -22.22 | 0.930 | 8 |
| Dense | 113 | 5.65 | 0.890 | -22.16 | 0.923 | 8 |
| Dense | 114 | 5.50 | 0.764 | -21.48 | 0.898 | 8 |
| Dense | 115 | 5.44 | 0.863 | -22.89 | 0.959 | 8 |
| Dense | 116 | 5.55 | 0.623 | -23.41 | 0.975 | 8 |
| Dense | 117 | 5.48 | 0.848 | -23.76 | 0.989 | 8 |
| Dense | 118 | 6.48 | 0.653 | -23.58 | 0.982 | 8 |
| Dense | 119 | 6.24 | 0.829 | -22.81 | 0.958 | 8 |
| NCP | 110 | 5.62 | 1.175 | -20.10 | 0.836 | 8 |
| NCP | 111 | 4.75 | 0.827 | -22.83 | 0.960 | 8 |
| NCP | 112 | 6.48 | 1.142 | -20.98 | 0.873 | 8 |
| NCP | 113 | 5.12 | 0.911 | -21.60 | 0.910 | 8 |
| NCP | 114 | 6.73 | 0.900 | -22.27 | 0.927 | 8 |
| NCP | 115 | 5.00 | 0.890 | -23.98 | 1.007 | 8 |
| NCP | 116 | 4.85 | 0.820 | -25.38 | 1.054 | 8 |
| NCP | 117 | 5.52 | 1.122 | -20.77 | 0.865 | 8 |
| NCP | 118 | 5.72 | 1.086 | -21.24 | 0.886 | 8 |
| NCP | 119 | 6.35 | 0.651 | -22.10 | 0.918 | 8 |
| FC | 110 | 4.22 | 0.758 | -22.54 | 0.942 | 8 |
| FC | 111 | 6.30 | 0.693 | -22.63 | 0.945 | 8 |
| FC | 112 | 5.44 | 0.835 | -23.24 | 0.974 | 8 |
| FC | 113 | 5.21 | 0.788 | -21.84 | 0.909 | 8 |
| FC | 114 | 6.70 | 0.832 | -21.96 | 0.910 | 8 |
| FC | 115 | 6.29 | 0.599 | -22.88 | 0.953 | 8 |
| FC | 116 | 6.04 | 0.682 | -22.91 | 0.954 | 8 |
| FC | 117 | 6.34 | 0.706 | -22.69 | 0.940 | 8 |
| FC | 118 | 7.07 | 0.969 | -24.10 | 1.004 | 8 |
| FC | 119 | 6.21 | 0.991 | -22.66 | 0.945 | 8 |
| ESN | 110 | 1.95 | 0.188 | -20.15 | 0.831 | 8 |
| ESN | 111 | 1.56 | 0.132 | -20.67 | 0.850 | 8 |
| ESN | 112 | 3.50 | 0.195 | -18.40 | 0.702 | 8 |
| ESN | 113 | 1.99 | 0.139 | -19.41 | 0.845 | 8 |
| ESN | 114 | 2.45 | 0.158 | -17.56 | 0.703 | 8 |
| ESN | 115 | 2.46 | 0.133 | -18.60 | 0.776 | 8 |
| ESN | 116 | 2.39 | 0.187 | -18.96 | 0.794 | 8 |
| ESN | 117 | 1.28 | 0.269 | -19.99 | 0.836 | 8 |
| ESN | 118 | 1.68 | 0.306 | -18.83 | 0.808 | 8 |
| ESN | 119 | 2.70 | 0.315 | -17.30 | 0.674 | 8 |
| RBF | 110 | 9.85 | 5.114 | -11.68 | 0.488 | 7 |
| RBF | 111 | 9.95 | 4.545 | -13.94 | 0.582 | 7 |
| RBF | 112 | 9.68 | 5.151 | -11.22 | 0.471 | 7 |
| RBF | 113 | 9.93 | 5.821 | -10.03 | 0.419 | 7 |
| RBF | 114 | 9.87 | 5.172 | -11.52 | 0.480 | 7 |
| RBF | 115 | 10.04 | 5.341 | -11.50 | 0.480 | 7 |
| RBF | 116 | 9.93 | 5.645 | -10.48 | 0.438 | 7 |
| RBF | 117 | 9.77 | 5.980 | -9.10 | 0.380 | 7 |
| RBF | 118 | 9.91 | 4.679 | -13.01 | 0.543 | 7 |
| RBF | 119 | 9.98 | 4.555 | -13.71 | 0.573 | 7 |
| RLS | 110 | 4.14 | 0.951 | -23.09 | 0.968 | 8 |
| RLS | 111 | 5.39 | 0.853 | -24.05 | 1.003 | 8 |
| RLS | 112 | 5.35 | 0.683 | -23.89 | 1.000 | 8 |
| RLS | 113 | 5.40 | 0.972 | -22.68 | 0.944 | 8 |
| RLS | 114 | 4.94 | 0.827 | -23.89 | 0.999 | 8 |
| RLS | 115 | 5.05 | 0.669 | -24.06 | 1.008 | 8 |
| RLS | 116 | 4.33 | 0.675 | -23.40 | 0.975 | 8 |
| RLS | 117 | 6.07 | 0.754 | -24.03 | 1.001 | 8 |
| RLS | 118 | 4.32 | 0.843 | -23.75 | 0.989 | 8 |
| RLS | 119 | 5.22 | 0.760 | -23.23 | 0.975 | 8 |

### B Auxiliary fields and sensitivity analyses

Table 4 reports results at *b* = −13 and *b* = 25 on the ten evaluation seeds, using the selected learning rates without retuning. At threshold 0.25 all runs pass the coarse decision, with the direction-count exceptions specified in the Results. Figure 7 shows the gain/rate sensitivity analysis, using only selection seeds 100–102. Among the 405 adaptive sensitivity outcomes reported here, 19 of the 135 reused selection-grid runs and 37 of the 270 new gain-scaled runs contain a non-finite cycle, leaving 349 finite points. Finite trajectories and A/B validity are separate checks.

**Table 4:**
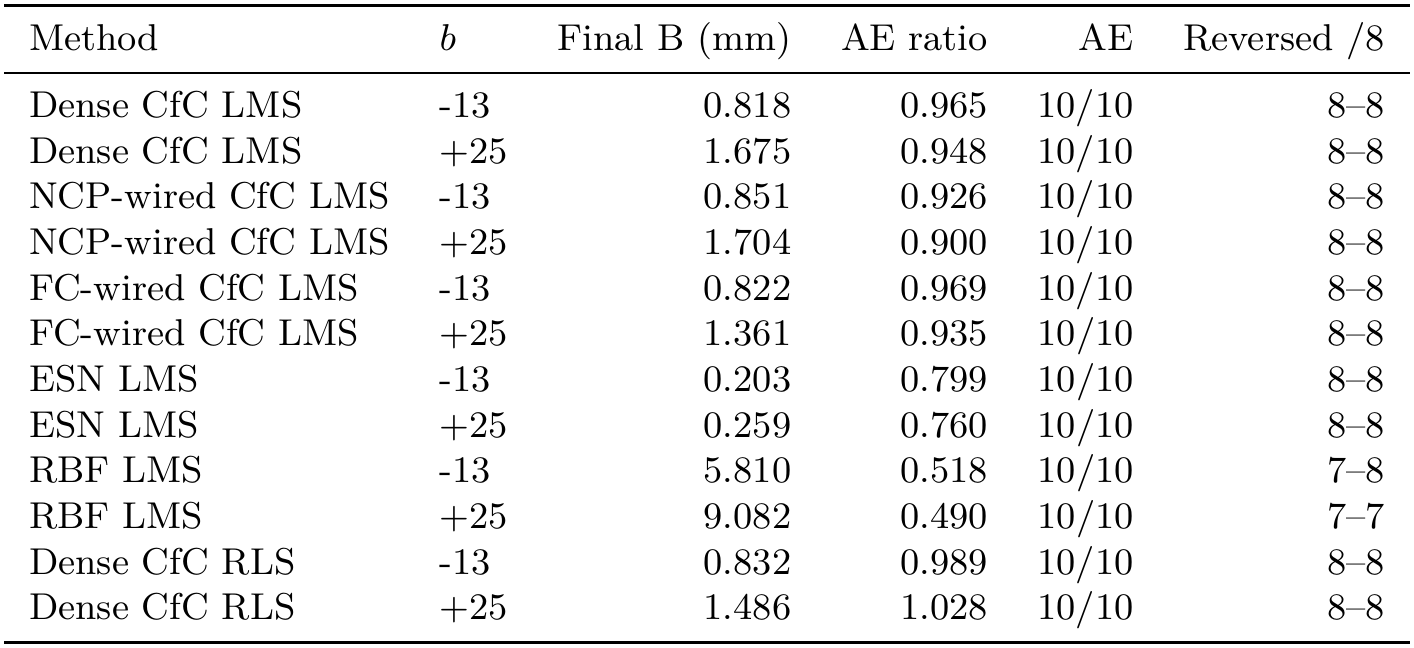
Reversed and stronger fields on evaluation seeds 110–119, without rate retuning. Final B and AE ratio are medians across ten seeds; AE counts seeds passing threshold 0.25; Reversed /8 gives the minimum– maximum direction count across seeds.

**Figure 7:**
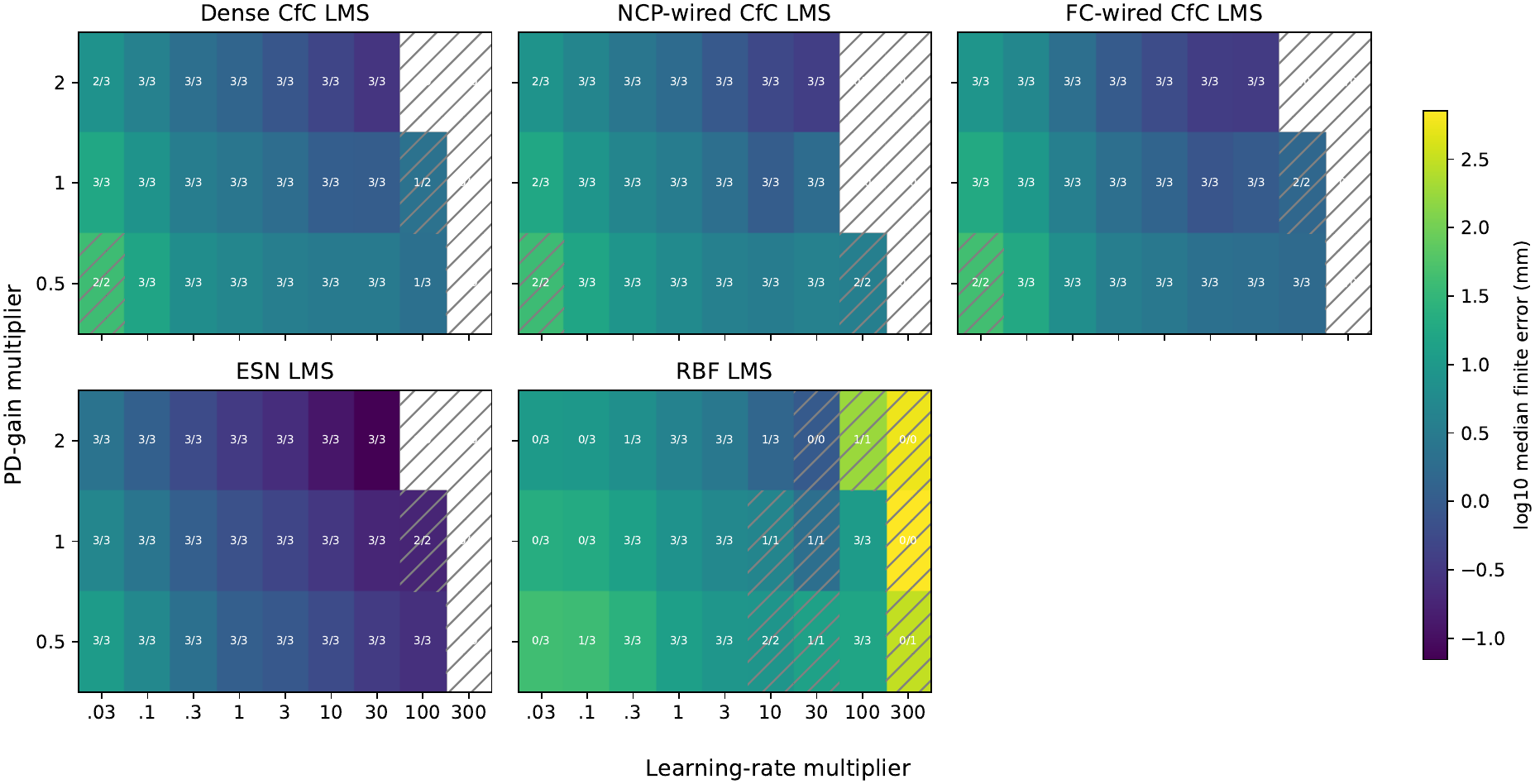
Selection-seed gain/rate sensitivity for the 400-epoch rerun. Colors show log_10_ median final error among finite runs. Cell text counts valid seeds passing the criterion over valid seeds; hatching marks at least one invalid seed. Empty colors indicate no finite run. Invalid and non-finite outcomes are retained in the machine-readable table. The selected rate is not changed after this analysis.

## C Reproducibility and provenance

### C.1 Reproduction materials and execution

The code and data archive described in the Data and code availability statement contains simulation code, model checkpoints, per-seed results, model-preparation and protocol settings, dataset manifests, and a dependency lockfile. The package’s `README`.md file provides instructions and commands for regenerating figures and tables, running diagnostics from saved readouts, repeating online experiments, and retraining offline models. Reproduction reports distinguish executed checks from available commands. File hashes verify the supplied code, models, and data before execution.

### C.2 Dataset provenance and validation

The dataset contains 65 models: 39 CfC checkpoints (three configurations, thirteen seeds), thirteen ESNs, and thirteen RBF models. Nine selection-seed CfC checkpoints were reused from a preparatory training run with identical settings; thirty evaluation-seed CfC models were newly trained. Initialization and wiring seeds, model and data hashes, and tensor/mask audits are recorded.

The supplied results comprise 843 protocol executions: 135 learning-rate selection runs, 333 evaluation runs (180 adaptive, 150 fixed-readout, and three deterministic PD-only), 315 new sensitivity runs, and 60 diagnostic reruns without washout. The 135 sensitivity entries at the original PD gains reuse selection runs rather than add executions. The diagnostics additionally include 120 field-on fixed-readout cycles (960 reaches), plus null-field fixed-readout cycles for reproduction checks. Non-finite selection and sensitivity outcomes are reported rather than silently removed.

The rerun also evaluated a direction-indexed torque-offset condition, which was omitted from this report after the outcomes were inspected.

For all 60 diagnostic reruns, cycle-level errors during the initial null-field, exposure, and catch phases match those of the corresponding evaluation runs. Readout weights remain unchanged during catch, reloading saved readouts reproduces their outputs, and common-input replay reproduces the recorded feedforward torques. The numerical source files and checkpoint tensors are unchanged by dataset packaging.

### C.3 NCP implementation details

The ncps 1.0.1 torch implementation stores connectivity masks as fixed arrays. A zero mask entry disables a connection in the corresponding weight matrix; a one entry allows its weight to be learned during offline training. For NCP-wired CfC, AutoNCP generates these masks from the group structure, sparsity setting, and wiring seed. The masks are not learned and are excluded from trainable parameter counts. They apply to the two candidate-output weight matrices, while within-group recurrent blocks remain dense and time-gating matrices are unmasked. Wiring-object synapse counts therefore do not count all effective connections. The supplied tensor/mask audit records the matrices, masks, and parameter counts used.

### C.4 Additional diagnostics on realized trajectories

Additional measurements evaluate the learned correction on the realized trajectories of the fixed pre- and post-exposure controllers. Here, the actor is the controller that generated a trajectory. We replay that trajectory’s inputs through both readouts to obtain their common-input output difference *δτ* (*x*_actor_), checking that the actor’s output reproduces its recorded feedforward torque. We evaluate Equation (7) by replacing *s*^*\**^ with the ideal compensation along the realized trajectory,

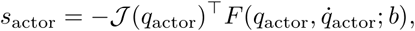

and recomputing the normalization denominator as *D*_actor_ = ⟨*s*_actor_, *s*_actor_⟩ . This compares torques at controller sample times, not an energy balance over RK4 substeps. Evaluating the other readout on these inputs gives a counterfactual output, not a counterfactual closed-loop trajectory, and does not separate the contribution of PD feedback. The data also report the root-mean-square magnitude of the feedback-torque vector.

## Declarations

### Competing interests

The author declares no competing interests.

### Funding

This research did not receive any specific grant from funding agencies in the public, commercial, or not-for-profit sectors.

### Author contributions

Jun Kobayashi: Conceptualization, Methodology, Software, Validation, Formal analysis, Investigation, Data curation, Visualization, Writing—original draft, Writing—review and editing.

### Data and code availability

The simulation code and experimental data are available in the GitHub release https://github.com/jkoba0512/cfc-online-adaptation/releases/tag/biorxiv-v4 and the versioned Zenodo archive https://doi.org/10.5281/zenodo.24094939. The archive includes the code, 65 model checkpoints, per-seed results, manifests, a dependency lockfile, and reproduction instructions. Validation reports document the checks performed and their scope. No human participant data were collected for this study.

### License

This preprint is distributed under the Creative Commons Attribution 4.0 International license (CC BY 4.0). The released code is provided under the MIT License identified in its LICENSE file.

## Footnotes

1 Official implementation: https://github.com/mlech26l/ncps.

## Notes

* OpenAI Codex and Anthropic Claude Code were used as assistive tools for planning, implementation support, manuscript drafting and editing, and consistency checks. Research decisions and interpretations were made by the author, who reviewed all content and takes responsibility for the work.

https://doi.org/10.5281/zenodo.24094939

https://github.com/jkoba0512/cfc-online-adaptation/releases/tag/biorxiv-v4

### Competing Interest Statement

The authors have declared no competing interest.

### Summary of Updates

This revision reframes the study around the distinction between catch-trial reversal, learned torque compensation, and performance after readout updates stop. It reports a new comparison of three CfC implementations, all trained for 400 epochs, with ESN, RBF, RLS, and fixed-readout baselines. Learning rates are selected on three model initializations and evaluated on ten separate initializations under three force fields. Compensation diagnostics and fixed-readout tests cover the same 60 main-field evaluation conditions. The diagnostic questions were motivated by earlier exploratory analyses, and their procedures were fixed before this rerun. The revised comparison uses the new evaluation data rather than earlier illustrative examples. Code and reproduction data are available through the GitHub biorxiv-v4 release and Zenodo DOI 10.5281/zenodo.24094939. The study does not claim a new learning algorithm, an isolated architecture advantage, or biological validation.

